# A Membrane Lipid Signature Unravels the Dynamic Landscape of Group 1 ILCs

**DOI:** 10.1101/2024.04.17.589821

**Authors:** Halle C. Frey, Xin Sun, Fatima Oudeif, Darleny L. Corona, Zijun He, Taejoon Won, Tracy L. Schultz, Vern B. Carruthers, Amale Laouar, Yasmina Laouar

## Abstract

In an era where the established lines between cell identities are blurred by intra-lineage plasticity, distinguishing between stable and transitional states becomes imperative. This challenge is particularly pronounced within the Group 1 ILC lineage, where the similarity and plasticity between NK cells and ILC1s obscure their classification and the assignment of their unique contributions to immune regulation. This study exploits the unique property of Asialo-GM1 (AsGM1)—a membrane lipid associated with cytotoxic attributes absent in ILC1s—as a definitive criterion to distinguish between these cells. By prioritizing cytotoxic potential as the cardinal differentiator, our strategic use of the AsGM1 signature achieved precise delineation of NK cells and ILC1s across tissues, validated by RNA-seq analysis. This capability extends beyond steady-state classifications, adeptly capturing the binary classification of NK cells and ILC1s during acute liver injury. By leveraging two established models of NK-to-ILC1 plasticity driven by TGFβ and *Toxoplasma gondii*, we demonstrate the stability of the AsGM1 signature, which sharply contrasts with the loss of Eomes. This signature identified a spectrum of known and novel NK cell derivatives—ILC1-like entities that bridge traditional binary classifications in aging and infection. The early detection of the AsGM1 signature at the immature NK (iNK) stage, preceding Eomes, and its stability, unaffected by transcriptional reprogramming that typically alters Eomes, position AsGM1 as a unique, site-agnostic marker for fate mapping NK-to-ILC1 plasticity. This provides a powerful tool to explore the expanding heterogeneity within the Group 1 ILC landscape, effectively transcending the ambiguity inherent to the NK-to-ILC1 continuum.

## INTRODUCTION

Innate lymphoid cells (ILCs) emerged as a dynamically evolving lineage within the immune system, playing a crucial role in orchestrating rapid responses to environmental stressors, microbial agents, and the complex cytokine microenvironment^1,2^. This broad activation mode positions ILCs as indispensable sentinels, finely tuned to react and adapt to environmental fluctuations^3–6^. Recent advances from transcriptomic and epigenomic studies have elucidated how ILCs adapt to these fluctuations, highlighting intra-lineage plasticity as a cornerstone mechanism^7,8^. This plasticity, evident across both human^9–15^ and murine studies^16–24^, introduces a significant layer of complexity to ILC classification, challenging the assignment of specific identities and roles. This challenge is particularly acute in the Group 1 ILC lineage, where significant plasticity and extensive overlapping traits between Natural Killer (NK) cells and Type 1 ILCs (ILC1s) obscure their precise identification, particularly in infection and cancer^19–25^.

The task of distinguishing ILC1s from NK cells relies on a suite of markers—Eomes and CD49b for NK cells, contrasted with CD200R, CD127, CD103, CD49a, and TRAIL for ILC1s depending on the anatomical site^26–29^. However, the efficacy of these markers in delineating ILC1s from NK cells is challenged by their variable expression. First, the presence or absence of Eomes is insufficient to distinguish NK cells from ILC1s due to its variable expression during NK cell activation^30^. Additionally, the low/absent Eomes expression in a subset of circulating human NK cells^31^ further questions its reliability as a discriminating marker. Second, CD200R can be upregulated on liver NK cells^21^ and is less effective in humans due to its expression in both ILC1s and NK cells^32^. Third, CD103 is inducible under the influence of TGFβ but is not expressed on most ILC1 subsets^19^.

Fourth, the expression of CD127 on immature NK cells^33^ (absent in mature NK cells) and its inconstant expression on ILC1s across tissues^34^ (e.g., absent in salivary gland ILC1s) makes this marker unreliable for distinguishing between NK cells and ILC1s. Fifth, the property of TRAIL as a selective marker for liver ILC1s is undermined by its inducible expression on NK cells in response to inflammatory cytokines^35,36^. Lastly, the upregulation of CD49a and down-regulation of CD49b on NK cells upon murine cytomegalovirus infection and cytokine stimulation highlights the challenges these markers present in identifying NK and ILC1 subsets^37–39^. Consequently, the current immunological framework lacks stable markers that can unambiguously distinguish ILC1s from NK cells across health and disease states. Moreover, recent data from fate mapping mouse models, adoptive transfers, and transcriptomic profiling in diverse biological contexts including endocrine tissues^19^, breast and prostate tumors^29^, and *Toxoplasma gondii* (*T. gondii*) infection^23^, identified a spectrum of intermediate subsets within the NK-to-ILC1 continuum that introduced an additional layer of complexity to the precise delineation of the Group 1 ILC landscape^19–25^. Defined by unique combinations of markers specific to either NK cells or ILC1s, the identification of these transitional states—known as ILC1-like cells—has disrupted the binary classification of ILC1s and NK cells, further complicating cell identity assignment within the Group 1 ILC landscape.

The quest for the precise segregation between ILC1s and NK cells stems from a key functional divergence^37,40^: the inherent cytotoxic capabilities of NK cells, crucial for the immune system’s primary defense against infections and tumors, are absent in ILC1s. In addressing this challenge, our strategy capitalized on the unique functional specificity of Asialo-GM1 (AsGM1)—a neutral membrane glycolipid^41–43^ known for its longstanding association with cytotoxic activities^44–48^. Through comprehensive analysis specific to each Group 1 ILC subset, complemented by transcriptomic profiling and models that foster intra-lineage plasticity, the strategic use of AsGM1 has not only enabled a precise demarcation of NK cells from their non-cytotoxic ILC1 counterparts across diverse anatomical sites, but also unveiled in environments conducive to NK-to-ILC1 plasticity a gradient of expression that captured a previously underappreciated level of diversity within the Group 1 ILC lineage. Our strategic focus on AsGM1, a membrane lipid marker empirically linked to cytotoxic activities, thus offers a unified approach to elucidate the diversity and dynamics of Group 1 ILCs, transcending the ambiguity inherent to the NK-to-ILC1 continuum.

## RESULTS

### AsGM1 strategically segregates Group 1 ILCs into ILC1 and NK cells across tissues

To evaluate the potential of AsGM1 in differentiating between the closely related Group 1 ILC subsets, we mapped its expression across a spectrum of anatomical sites—including lymphoid (bone marrow [BM], mesenteric lymph nodes [mLN], and spleen), circulatory (blood), non-lymphoid (liver), and mucosal (lung, and small intestine lamina propria [siLP]) sites (**Fig. 1** and **Supplementary** Fig. 1**, 2**). Analysis across these sites demonstrated a clear demarcation of AsGM1 expression, effectively segregating Group 1 ILCs into two well-defined subsets: AsGM1^-^ and AsGM1^+^ (**Fig. 1a**). AsGM1^+^ ILCs were predominantly found in the Group 1 ILC niches of the spleen, BM, blood, and lungs, while AsGM1^-^ ILCs were more abundant in the liver and mLN, and distinctly predominant in the siLP (**Supplementary** Fig. 1**, 2a**). This distribution pattern was validated by the anti-AsGM1 antibody treatment, which resulted in a significant depletion of the splenic Group 1 ILCs, in contrast to a modest decrease in the siLP which harbors a significant population of AsGM1^-^ cells (**Supplementary** Fig. 2a, c). These observations, further supported by the overlap in AsGM1^-^ ILC and ILC1 frequencies (**Fig. 1e**), underscore a parallel between the anatomical distribution of AsGM1-segregated ILCs and that of ILC1s and NK cells^34^. Subsequent analysis contrasting the expression profiles of AsGM1-segregated ILCs with the known marker signatures of ILC1s and NK cells, also revealed a significant parallel (**Fig. 1b-d** and **Supplementary** Fig. 2b). Precisely, AsGM1^-^ ILCs across examined sites displayed an ILC1 profile characterized by the expression of CD200R and T-bet (with liver ILCs additionally expressing TRAIL) while lacking Eomes and CD49b. Conversely, AsGM1^+^ ILCs exhibited traits of NK cells, marked by the expression of Eomes, T-bet, and CD49b and the lack of CD200R and TRAIL (**Fig. 1b, c**). AsGM1^+^ ILCs also displayed a wide array of activating and inhibitory markers including Ly49C/I, Ly49D, Ly49G2, Ly49H, NKG2ACE, KLRG1, and CD94. In contrast, AsGM1^-^ ILCs were devoid of this NK cell signature, conclusively aligning them with the characteristic profile of ILC1s (**Fig. 1d**).

**Figure 1.**
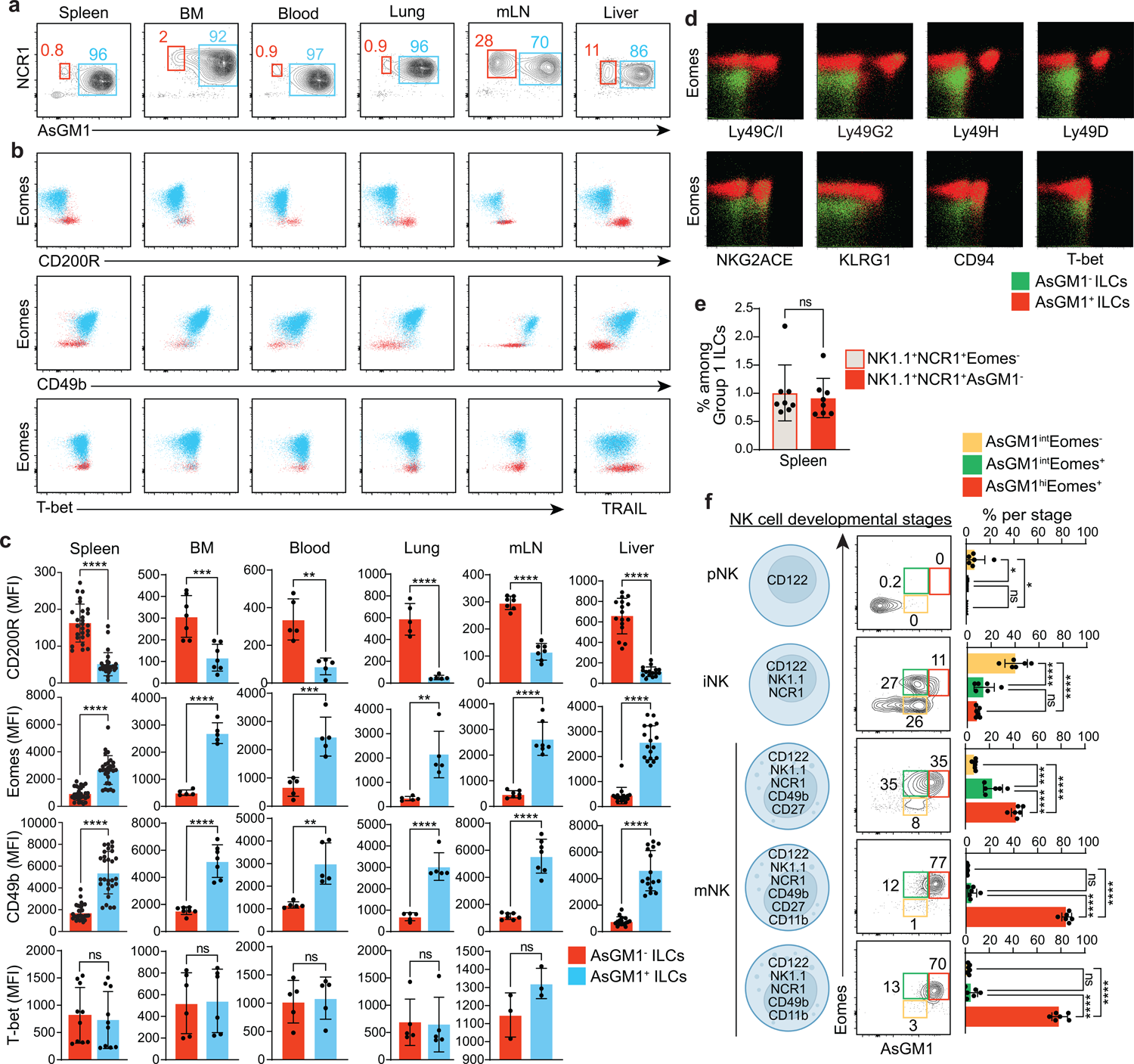
Mapping AsGM1 expression across Group 1 ILCs. (**a**) Distribution of NCR1 vs. AsGM1 in CD3^-^NK1.1^+^ ILCs in the spleen, bone marrow (BM), blood, lung, mesenteric lymph nodes (mLN), and liver. (**b**) Profiles of Eomes, CD200R, CD49b, T-bet, and TRAIL in AsGM1^-^ and AsGM1^+^ ILCs across tissues. (**c**) Mean Fluorescence Intensity (MFI) for CD200R, Eomes, CD49b, and T-bet in AsGM1^-^ and AsGM1^+^ ILCs across tissues. (**d**) Comparison of activating and inhibitory NK cell markers in AsGM1^-^ and AsGM1^+^ ILCs in the spleen. (**e**) Proportion of Group 1 ILCs delineated as CD3^-^NK1.1^+^NCR1^+^Eomes^-^ vs. CD3^-^NK1.1^+^NCR1^+^AsGM1^-^ in the spleen. (**f**) Distribution of AsGM1 vs. Eomes expression across five stages of NK cell development including precursor NK (pNK), immature NK (iNK), and CD11b^-^CD27^+^, CD11b^+^CD27^+^, and CD11b^+^CD27^-^ mature NK (mNK) stages. Frequencies of AsGM1^int^Eomes^-^, AsGM1^int^Eomes^+^ and AsGM1^hi^Eomes^+^ subsets within each developmental stage. Data represent 3 independent experiments: spleen (*n* = 27), BM (*n* = 6), blood (*n* = 5), lung (*n* = 5), mLN (*n* = 7), and liver (*n* = 16). Statistical validation employed unpaired t-tests (**c, e**) and one-way ANOVA (**f**), with significance indicated by \**p*<0.05, \*\**p*<0.01, \*\*\**p*<0.001, \*\*\*\**p*<0.0001. Error bars represent mean ± s.d. Refer to Supplementary Figures 1 and 2 for additional information.

To determine whether AsGM1 signature in NK cells occurs at a mature stage or at a precursor level, we mapped its expression across five stages of NK cell development (**Fig. 1f**). Results traced AsGM1 expression to early development, first detected at the immature NK (iNK) stage (**Fig. 1f**)—a crucial phase characterized by the initial detection of Eomes^49,50^ and inhibitory/activating receptors^51,52^, essential for functional maturation. Although Eomes and AsGM1 first appear at the iNK stage, our data indicates that AsGM1 expression precedes that of Eomes (**Fig. 1f**). This sequential expression pattern at the iNK stage—which is known to foster immature NK cells as well as ILC1 precursors^53,54^—underscores a critical juncture where AsGM1^+^ cells become likely poised to diverge into the NK cell lineage, with the subsequent Eomes expression further driving NK cell lineage specification^49,50^. Collectively, these data demonstrate the remarkable specificity of AsGM1 in delineating NK and ILC1 identities at a mature stage and segregating their developmental trajectories at an immature level.

### Functional and transcriptomic characterization of AsGM1-segregated ILCs

To conclusively establish the identity of AsGM1-segregated ILCs as bona fide ILC1s and NK cells, we used RNA-sequencing to compare their transcriptomic profiles (**Fig. 2a-d**). Using *Rag2^-/-^* mice, selected for their enriched ILC populations resulting from the absence of adaptive immune cells, enabled precise isolation of AsGM1^-^ and AsGM1^+^ subsets within Group 1 ILCs in the liver, where the frequency of both subsets is notably high^55^ (**Fig. 2a** and **Supplementary** Fig. 3). The comparison of transcriptomic profiles between AsGM1^-^ and AsGM1^+^ subsets revealed a signature of 281 differentially expressed genes (DEGs) (**Fig. 2b**). Among these DEGs, genes traditionally associated with ILC1s^56^, such as *Cd200r1*, *Cd200r2*, *Cxcr6*, *Il7r*, *Itga1*, *Tmem176a*, *Tmem176b*, and *Tnfsf10*, were predominantly expressed in AsGM1^-^ ILCs but notably lacking in AsGM1^+^ ILCs. Conversely, genes characteristic of NK cells^56^, including *Eomes*, *Prf1*, *Irf8*, *Gzmk*, *Klra4*, *Klra7*, and *Klra8*, were abundantly expressed in AsGM1^+^ ILCs but markedly reduced in the AsGM1^-^ subset (**Fig. 2c, d**). To validate these findings, we cross-referenced our data with transcriptomic datasets from independent studies that delineated ILC1s and NK cells in the liver, spleen, and intestine by distinct expression patterns of Eomes, TRAIL, or CD127^37,57^ (**Fig. 2e-g**). Across all heatmaps, our analysis revealed a uniform pattern of differential gene expression, firmly establishing the identity of AsGM1^-^ and AsGM1^+^ Group 1 ILCs as authentic ILC1 and NK cell subsets, respectively.

**Figure 2.**
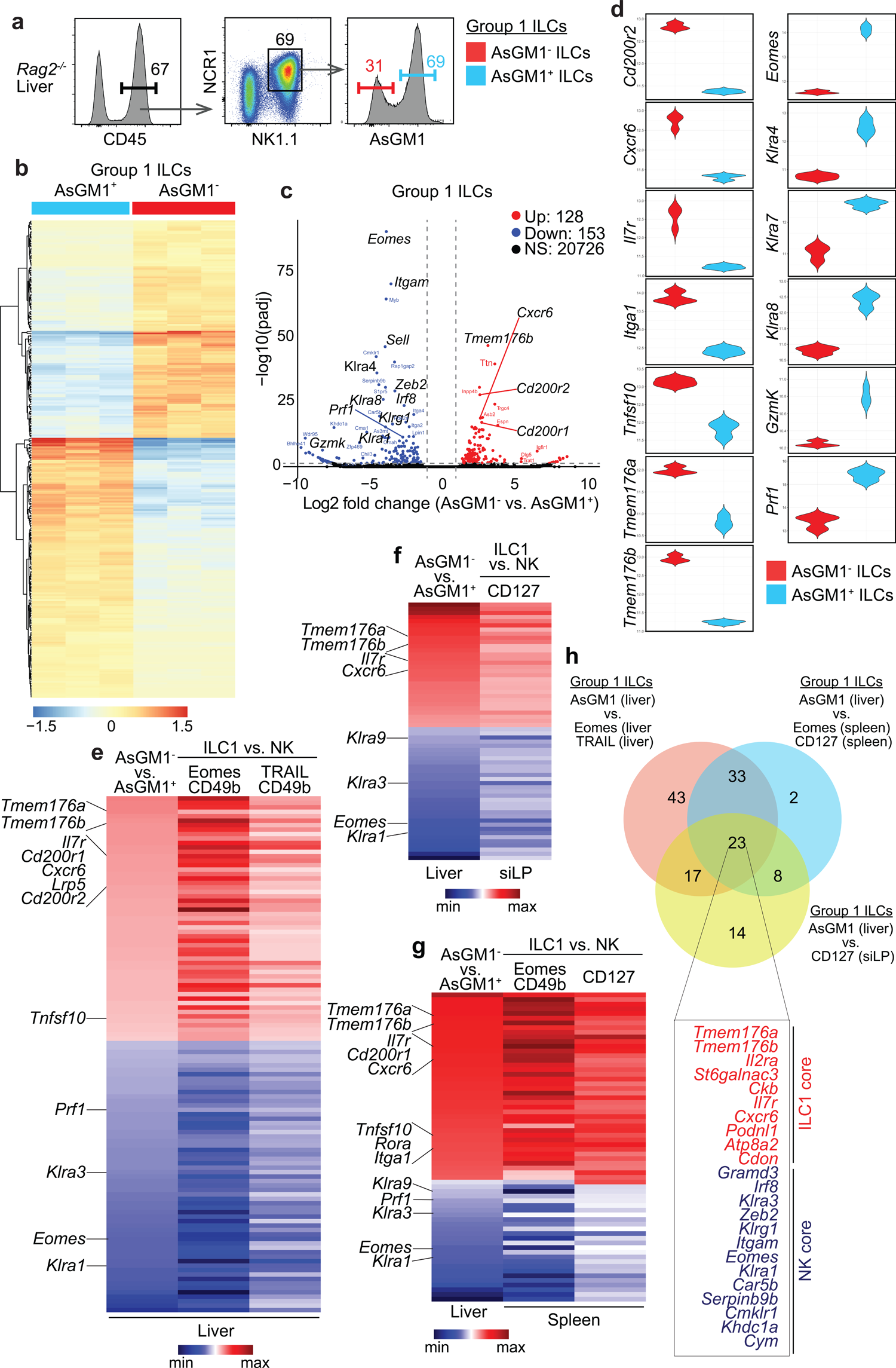
Profiles of AsGM1^-^ and AsGM1^+^ Group 1 ILCs match ILC1s and NK cells. (**a-d**) Transcriptomic profiling of AsGM1-segregated Group 1 ILCs. (**a**) Gating strategy for sorting AsGM1^-^ and AsGM1^+^ Group 1 ILCs from the liver of *Rag2^-/-^* mice. (**b**) Heatmap showing the differential gene expression between AsGM1^-^ and AsGM1^+^ Group 1 ILCs. (**c**) Volcano plot contrasting gene expression between AsGM1^-^ and AsGM1^+^ Group 1 ILCs, highlighting differentially expressed genes (DEGs) based on significance (*p*-values < 0.05) and magnitude of change (fold-change > 1.5): up-regulated (red), down-regulated (blue), and non-significant changes (black). (**d**) Violin plots illustrating the expression patterns of DEGs associated with ILC1 (left) or NK cell (right) profiles. (**e-g**) Comparative analysis of DEGs from AsGM1-segregated Group 1 ILCs [log_2_(AsGM1^-^/AsGM1^+^)] against DEGs from existing datasets of ILC1 and NK cell subsets [log_2_(ILC1/NK)], identified based on marker combinations of Eomes and CD49b, TRAIL and CD49b, or CD127 across the liver^37^ (**e**), intestine^57^ (**f**), and the spleen^37,57^ (**g**). (**h**) Venn diagram illustrating the overlap in gene expression across Group 1 ILC profiles, distinguished by AsGM1 in the liver versus ILC1s and NK cells the liver^37^, spleen^37,57^, and intestine^57^. A list of 23 genes at the intersection of all comparison is provided. Data representative of *n* = 3 show individual mice in (**b**) and mean in (**c-h**). Refer to Supplementary Figure 3 for additional information.

Contrasting AsGM1-defined gene expression in the liver with 5 profiles identified by Eomes or TRAIL in the liver^37^, Eomes or CD127 in the spleen^37,57^, and CD127 in the intestine^57^, via Venn diagram analysis, revealed a consensus set of 23 genes, comprising 10 ILC1 core genes and 13 NK core genes. (**Fig. 2h**). Within the intersecting ILC1 core, *Il7r*^57^ and *Ckb*^60^, known to be expressed in ILC progenitors (ILCPs)^58^, supports the unique origin of ILC1s from ILCPs, diverging from the NK cell lineage. *Podnl1*^59,60^, integral to the ILC1 core for cell adhesion and cellular matrix interactions^61^, aligns with the predominant localization of ILC1s as tissue-residents—as opposed to the circulatory nature of NK cells. *St6galnac3*, a gene that encodes a sialyltransferase responsible for attaching sialic acid to glycoproteins and glycolipids^62^, contrasts sharply with the sialic acid-deficient AsGM1 glycoprotein^63^. Accordingly, the ubiquitous expression of *St6galnac3*^60^ in ILC1s outlines a distinct sialylation landscape that diverges ILC1s from the cytotoxic profile of NK cells, which is linked to AsGM1^44–48^. Within the intersecting NK core, *Gramd3* expression aligns with its association with optimal cytotoxicity mediated by DNAM^64,65^. *Serpinb9b*, a gene encoding a serine protease inhibitor, plays a key role in immunoregulation by obstructing Granzyme B activity, ensuring that NK cells execute targeted cytotoxic actions without self-inflicted damage^66^. Its identification in our intersecting NK core^49,57,60^, aligns with its pivotal role in protecting the cytotoxic function of NK cells, absent in ILC1s. *Cmklr1*, a chemokine receptor associated with the migration of human and murine NK cells to sites of tumors^67–69^, emerged as a gene signature of NK cells^68^, analogous to the integral *Cxcr6* expression in ILC1s. Lastly, it is unsurprising that *Itgam*^56,60^, *Khdc1a*^60^, and *Cym*^60^ genes emerged in our intersecting NK core, given their association with T-bet and/or Eomes^49^.

To conclusively establish functional parallels—between AsGM1^-^ ILCs and ILC1s on one hand, and AsGM1^+^ ILCs and NK cells on the other—we examined the cytokine responses of both AsGM1^-^ and AsGM1^+^ Group 1 ILCs to IL-12 and IL-18, which induce IFNγ production^70^, and IL-12 plus IL-15, which triggers Granzyme B (GzmB) expression^71^. To this end, we used the siLP—a mucosal site enriched with NK cells, ILC1s, and ILC3s— which provides an optimal context for comparing AsGM1^-^ and AsGM1^+^ Group 1 ILCs, while also allowing ILC3s to serve as an internal control (**Fig. 3a-c**). While both AsGM1^-^ and AsGM1^+^ Group 1 ILCs exhibited comparable IFNγ output, underscoring a functional overlap between AsGM1^-^ and AsGM1^+^ subsets, the cytotoxic function reflected by GzmB expression was specific to the AsGM1^+^ subset (**Fig. 3d**). This distinction emphasizes the functional divergence within AsGM1-segregated ILCs, mirroring the functional dichotomy between ILC1s and NK cell subsets^37,40^. These findings, stemming from integrated transcriptomic profiling and functional assays, introduce a new framework that adeptly distinguishes NK cells from ILC1s, via a single-marker approach predicated on cytotoxic potential.

**Figure 3.**
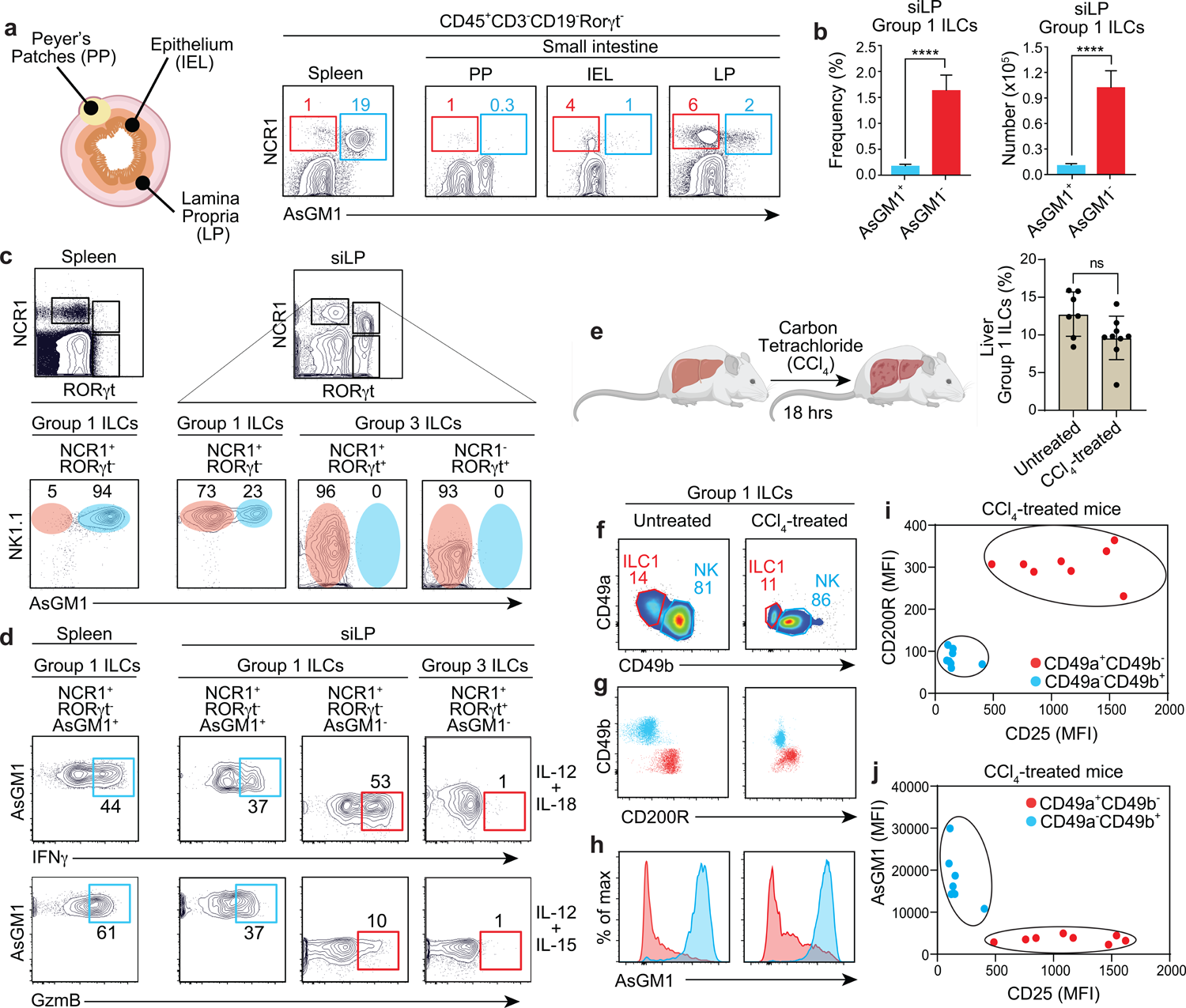
Stable AsGM1 signatures define Group 1 ILCs from different anatomical locations upon acute activation. Patterns of AsGM1 expression in Group 1 ILCs from the intestine (**a-d**) and liver (**e-j**), subjected to *in vitro* and *in vivo* stimulations, respectively. (**a**) Pattern of AsGM1 expression within Group 1 ILCs of the spleen and small intestine Peyer’s patches (PP), intraepithelial lymphocytes (IEL), and lamina propria (siLP). (**b**) Frequency and numbers of AsGM1-segregated Group 1 ILCs within the siLP. (**c**) AsGM1 expression across spleen Group 1 ILCs (NCR1^+^Rorγt^-^), siLP Group 1 ILCs (NCR1^+^Rorγt^-^), and siLP Group 3 ILCs (NCR1^+^Rorγt^+^ and NCR1^-^Rorγt^+^). (**d**) IFNγ and GzmB expression in AsGM1^-^ and AsGM1^+^ Group 1 ILCs of the siLP in response to stimulation with IL-12 plus IL-18 or IL-12 plus IL-15. Spleen NK cells and siLP ILC3s are used as controls. (**e**) Frequency of hepatic Group 1 ILCs before and 18 hours after CCl_4_ treatment. (**f**) Distribution of CD49a vs. CD49b within hepatic Group 1 ILCs before and after CCl_4_ treatment. (**g**) Overlay comparison of CD49b vs. CD200R distribution between ILC1s and NK cells before and after CCl_4_ treatment. (**h**) Patterns of AsGM1 expression in ILC1s and NK cells before and after CCl_4_ treatment. (**i, j**) Mean Fluorescence Intensity (MFI) of CD200R vs. CD25 (**i**) and AsGM1 vs. CD25 (**j**) in ILC1s and NK cells after CCl_4_ exposure. Graphs show MFI values from individual mice. Data represent 3 (**a-d**) and 2 (**e-j**) independent experiments, with *n* = 3-6 (**a-d**) and *n* = 7-8 (**e-j**). Statistical validation employed unpaired t-test (**b, e**), with ****p<0.0001 indicating significance. Error bars represent mean ± s.d.

### Stable AsGM1 expression in the continuum from homeostasis to acute response

The task of accurately distinguishing ILC1s from NK cells becomes increasingly challenging under physiological stress, where dynamic shifts in marker expression often occur^19,21,30,35–37,72^. To evaluate the effectiveness of AsGM1 in resolving the heterogeneity of Group 1 ILCs under physiological stress or injury, we used a carbon tetrachloride (CCl_4_)-induced model of acute liver injury, known to trigger acute activation of ILC1s in the liver^73^. Following established protocols^73^, C57BL/6 mice received an intraperitoneal injection of CCl_4_; after 18 hours, hepatic Group 1 ILCs were analyzed to compare AsGM1 expression in steady state versus acute activation (**Fig. 3e-j**). We applied established markers to differentiate ILC1s (CD3^-^NK1.1^+^CD49a^+^CD49b^-^CD200R^+^) from NK cells (CD3^-^NK1.1^+^CD49a^-^CD49b^+^CD200R^-^) (**Fig. 3f, g**). Additionally, we used CD25 as an indicator of activation to verify the immediate response of liver ILC1s to CCl4 treatment^73^ (**Fig. 3i**). Using this experimental setup, we proceeded to assess the patterns of AsGM1 expression in response to acute activation. Our data showed that AsGM1 expression within Group 1 ILCs remained a reliable marker for distinguishing ILC1s from NK cells in CCl_4_-treated mice compared to control counterparts (**Fig. 3h**). Specifically, ILC1s—whether in a resting state or under acute activation—maintained an AsGM1^-^ profile, while NK cells invariably showed an AsGM1^+^ profile (**Fig. 3h, j**). This remarkable stability, despite the physiological stress triggered by acute liver injury, underscored AsGM1’s unparalleled precision in delineating the binary classification of NK cells and ILC1s, bridging the continuum from steady state to immediate response with a clear demarcation.

### AsGM1 as a unique tool for tracing intra-lineage plasticity within Group 1 ILCs

An inherent challenge posed by cellular plasticity lies in the transcriptomic reprogramming it entails, which modifies cellular identities making it difficult to ascertain which cells are stable and which are plastic^74^. We posit that AsGM1 is uniquely qualified to track NK-to-ILC1 plasticity, aiming to delineate ILC1-like cells to their NK cell origin. This hypothesis rests on the premise that AsGM1, a cellular membrane lipid component^41,42^, remains unaffected by the transcriptomic reprogramming that governs NK-to-ILC1 plasticity, potentially serving as a stable indicator of plasticity. To this end, we used the gold standard *in vivo* model of NK-to-ILC1 plasticity driven by TGFβ in the salivary glands (SG)^19^. In line with these studies^19^, our findings confirmed a transition from a primarily NK cell phenotype (CD49a^-^CD49b^+^) in the SGs of CD11c^dnR^ mice^75,76^ (characterized by TGFβ-resistant NK cells) to a dominant ILC1-like profile (CD49a^+^CD49b^+^, known as SG ILCs) in wild-type SGs (TGFβ-responsive NK cells) (**Fig. 4a**). During this transition, Eomes expression was downregulated, as evidenced by intermediate levels in SG ILCs. In contrast, AsGM1 expression remained stable, enabling accurate tracking of NK-to-ILC1 plasticity (**Fig. 4b, c**). To further validate the stability of AsGM1, we used a well-established *in vitro* model of NK-to-ILC1 plasticity, which involves treating splenic NK cells with TGFβ plus IL-15 to induce their conversion into ILC1-like entities^20^ (**Fig. 4d, e**). After a seven-day exposure to this cytokine milieu, the resultant cells exhibited a downregulation of Eomes along with an upregulation of CD49a, CD103, and TRAIL, mirroring outcomes of NK cell conversion associated with TGFβ in prior studies^19,20^. Importantly, AsGM1 expression remained constant throughout this transition, underscoring its stability during NK-to-ILC1 plasticity (**Fig. 4d, e**).

**Figure 4.**
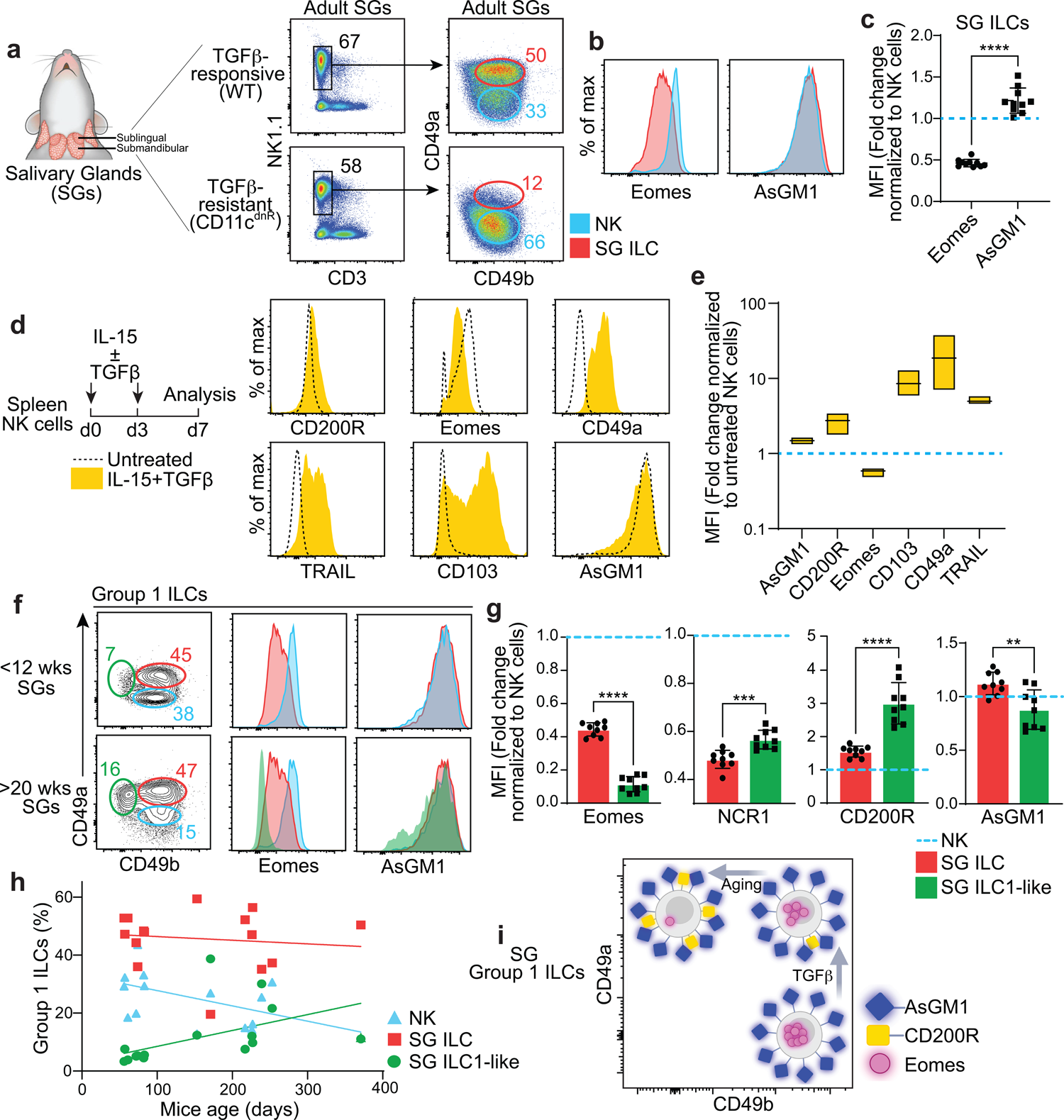
Tracing NK-to-ILC1 plasticity in the salivary glands via AsGM1. (**a**) Distribution of CD49a vs. CD49b among Group 1 ILCs (CD45^+^CD3^-^NK1.1^+^) in the salivary glands (SGs) of adult wild-type (WT) and CD11c^dnR^ (TGFβR-resistant NK cells) mice. (**b**) Expression of Eomes and AsGM1 in NK cells and SG ILCs from wild-type SGs. (**c**) MFI of Eomes and AsGM1 expression in SG ILCs relative normalized to NK cells in wild-type SGs. (**d**) Expression of CD200R, Eomes, CD49a, TRAIL, CD103, and AsGM1 in splenic NK cells before (dotted line) and after IL-15 plus TGFβ treatment (yellow). (**e**) MFI of AsGM1, CD200R, Eomes, CD103, CD49a, and TRAIL under IL-15 plus TGFβ treatment normalized to untreated controls. (**f**) Distribution of CD49a vs. CD49b in SG Group 1 ILCs across adult (2-4 months) and aged (5-12 months) wild-type mice. Gates indicate NK cells (blue), SG ILCs (red), and SG-ILC1-like cells (green). Histograms show expression of Eomes and AsGM1 among NK cells, SG ILCs, and SG ILC1-like cells. (**g**) MFI of Eomes, NCR1, CD200R, and AsGM1 in SG ILCs and SG ILC1-like cells normalized to NK cells in aged (5-12 months) wild-type mice. (**h**) Age-related frequency shifts in NK cells, SG ILCs, and SG ILC1-like cell subsets. (**i**) Schematic representation illustrating the stable expression of AsGM1 against the fluctuating expression of CD200R and Eomes in SG Group 1 ILCs, highlighting an age-associated continuum. Data represent 3 independent experiments: *n* = 11 (**a-c**), *n* = 3 (**d-e**), *n* = 13 (**f-h**) for wild-type mice, with *n* = 5 for CD11c^dnR^ mice (**a**). Statistical validation employed unpaired t-test (**c, g**) and one-way ANOVA (**e**), with significance indicated by \*\**p*<0.01, \*\*\**p*<0.001, \*\*\*\**p*<0.0001. Error bars represent mean ± s.d.

Given that aging introduces qualitative alterations in the microenvironment^77^, we focused on how these shifts affect NK-to-ILC1 plasticity within the aging SGs, postulating that the transformed milieu of an aged setting might lead to distinct plasticity outcomes. Within this unique context, AsGM1 demonstrated its potential as a tracing marker by capturing a previously undescribed age-dependent ILC1-like state (AsGM1^+^CD200R^+^ Eomes^-^CD49a^+^CD49b^-^), designated as SG ILC1-like cells (**Fig. 4f-h**). This finding extends the known spectrum of SG Group 1 ILC diversity to incorporate NK cells, SG ILCs, and the age-dependent SG ILC1-like cells, unified by their shared AsGM1 expression (**Fig. 4f**). The distinct gradients established by Eomes and CD200R expression among these three subsets, highlight an age-associated continuum of plasticity within the Group 1 ILC lineage, evolving from NK cells to SG ILCs, and ultimately to SG ILC1-like cells. This progression, characterized by the reciprocal modulation of Eomes and CD200R together with stable AsGM1 expression, places SG ILC1-like cells at an advanced stage in the NK-to-ILC1 continuum, influenced by the unique aging microenvironment (**Fig. 4g**). Additionally, age-related shifts in cell prevalence, highlighted by a decrease in NK cells and an increase in SG ILC1-like cells (**Fig. 4h**), illustrate an age-dependent adaptation and reconfiguration of the SG Group 1 ILC landscape, distinctly captured by AsGM1 tracing (**Fig. 4i**). This continuum depicted in aged SGs strikingly mirrors the transition from NK cells (CD49b^-^CD49a^-^) to intermediate ILC1s (CD49b^+^CD49a^+^) and to ILC1s (CD49b^-^CD49a^+^) in the tumor microenvironment^20^, hinting at a unified adaptation strategy of Group 1 ILCs in aging and cancer, two different scenarios united by gradual timelines of cellular and molecular damage accumulation.

### AsGM1 gradient reveals greater spectrum of Group 1 ILC diversity during *T. gondii* infection

To assess the potential broad application of AsGM1 tracing across other immunological contexts, we employed *T. gondii* infection which represents a distinct model of NK-to-ILC1 plasticity to evaluate AsGM1 tracing in the specific context of infection-driven plasticity^23^.

Following established protocols, we inoculated C57BL/6 mice with *T. gondii* and assessed alterations within the Group 1 ILC lineage in the spleen, lung, and liver 14 days post-infection^23^ (**Fig. 5**). Quantitative analysis revealed an expansion of the ILC1 pool and a concurrent reduction of the NK cell pool post-infection, consistent with NK-to-ILC1 plasticity^23^ (**Fig. 5a, b**). Additional analysis showing an upregulation of Ly6C on ILC1s, a key indicator of plasticity^23^, further substantiates this shift (**Fig. 5c, e** and **Supplementary** Fig. 4). Further scrutiny of the liver ILC1 pool (NK1.1^+^NCR1^+^CD49a^+^Eomes^-^) from *T. gondii*-infected mice revealed a peculiar subset with NK cell characteristics, including the lack of CD200R and the expression of CD49b (**Fig. 5d**). This subset, displaying an atypical ILC1 profile characterized by NK1.1^+^NCR1^+^CD49a^+^Eomes^-^CD200R^-^CD49b^+^ phenotype, also expressed AsGM1^int^, hinting at a transitional state from an NK cell origin (**Fig. 5f**). Mapping AsGM1 expression across Group 1 ILCs has thus unveiled a distinct gradient, capturing three subsets 14 days post-infection: conventional NK cells (AsGM1^+^), classical ILC1s (AsGM1^-^), and, notably, transitional ILC1-like cells^23^, which we identified by their intermediate AsGM1 expression (AsGM1^int^) (**Fig. 5g**).

**Figure 5.**
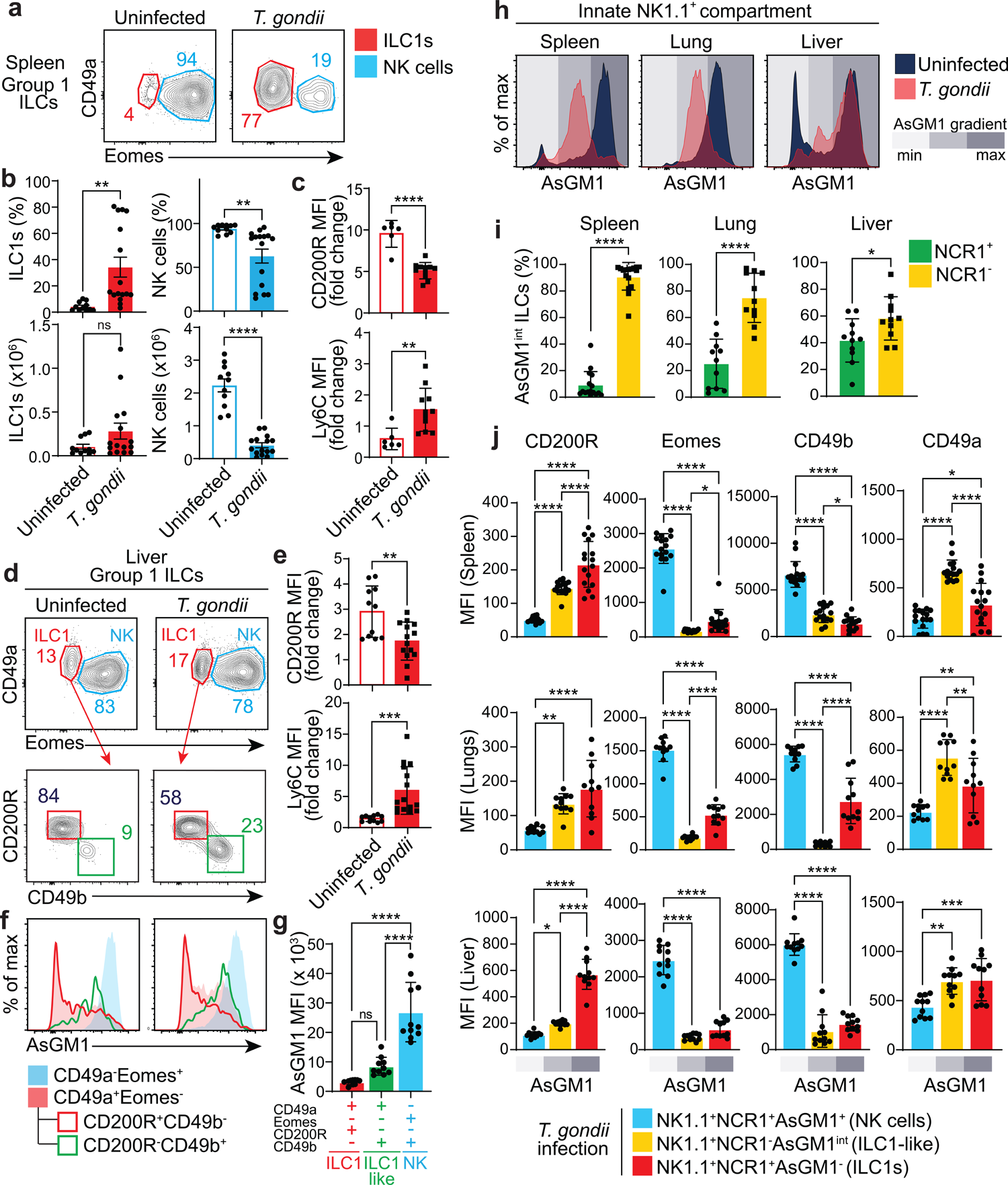
AsGM1 gradient driven by *T. gondii* infection captures two distinct ILC1-like subsets. (**a-j**) Analysis of uninfected and *T. gondii*-infected mice 14 days post-infection. (**a**) Distribution of CD49a vs. Eomes among splenic Group 1 ILCs (CD45^+^CD3^-^ NK1.1^+^NCR1^+^) from uninfected and *T. gondii*-infected mice. (**b**) Frequency and numbers of ILC1s and NK cells among splenic Group 1 ILCs from uninfected and *T. gondii*-infected mice. (**c**) MFI of CD200R and Ly6C in splenic ILC1s normalized to splenic NK cells from uninfected and *T. gondii*-infected mice. (**d**) Distribution of CD49a vs. Eomes among liver Group 1 ILCs from uninfected and *T. gondii*-infected mice. Distribution of CD200R vs. CD49b among liver ILC1s from uninfected and *T. gondii*-infected mice. (**e**) MFI of CD200R and Ly6C in liver ILC1s normalized to liver NK cells of uninfected and *T. gondii*-infected mice. (**f**) AsGM1 expression across liver Group 1 ILC subsets from uninfected and *T. gondii*-infected mice. (**g**) MFI of AsGM1 in ILC1s, ILC1-like cells, and NK cells in the liver of *T. gondii*-infected mice. (**h**) AsGM1 expression in CD45^+^CD3^-^NK1.1^+^ ILCs from the spleen, lung, and liver of uninfected and *T. gondii*-infected mice. A graded grey bar indicates the gradient of AsGM1 expression. (**i**) Distribution of NCR1^+^ and NCR1^-^ ILC subsets among NK1.1^+^AsGM1^int^ ILCs across spleen, lung, and liver in *T. gondii*-infected mice. (**j**) MFI of CD200R, Eomes, CD49b, and CD49a in NK1.1^+^NCR1^-^AsGM1^int^ ILCs compared with NK cells and ILC1s in the spleen, lung, and liver of *T. gondii*-infected mice. A graded grey bar indicates AsGM1 expression. Data represent 2 independent experiments, with *n* = 6-11 uninfected mice and *n* = 11-16 infected mice. Statistical validation employed unpaired t-test (**b, c, e, i**) and one-way ANOVA (**g, j**), with significance indicated by \**p*<0.05, \*\**p*<0.01, \*\*\**p*<0.001, \*\*\*\**p*<0.0001. Error bars represent mean ± s.e.m. (**b**) and mean ± s.d. (**c, e, g, i, j**).

An additional subset of AsGM1^int^ ILC1-like cells, distinguished by the lack of NCR1, was identified within the Group 1 ILC population across all sampled tissues, with a notable predominance in the infected spleens and lungs (**Fig. 5h, i** and **Supplementary** Fig. 5a). This distinct subset displayed a peculiar phenotype, characterized by NK1.1 and CD49a expression, lack of NCR1, CD49b, and Eomes, and the expression of intermediate levels of CD200R and AsGM1, consistent with a transitional stage in the NK-to-ILC1 continuum driven by *T. gondii* infection (**Fig. 5j**). This is further substantiated by infection-associated shifts in cell prevalence, marked by a reduction in NK cells and a rise in AsGM1^int^NCR1^-^ILC1-like cells (**Supplementary** Fig. 5b, c), demonstrating a significant, infection-driven reconfiguration of the Group 1 ILC landscape. This newly characterized population, which is likely overlooked due to the lack of NCR1, introduces a fourth subset to the Group 1 ILC spectrum, expanding its diversity to include two stable (AsGM1^-^ ILC1s and AsGM1^+^ NK cells) subsets and two infection-driven ILC1-like states (AsGM1^int^NCR1^+^ and AsGM1^int^NCR1^-^), all distinctly captured by the AsGM1 expression gradient.

## DISCUSSION

The evolution from a monolithic view of NK cells to a diverse framework—Group 1 ILCs— marked a pivotal shift in our understanding of the innate lymphoid system’s architecture^78^. Initially, ILC1s were conflated with NK cells due to shared core markers—NK1.1, NCR1, T-bet, and IFNγ—that blurred their unique identities^25^. The recognition of Group 1 ILCs as a diverse lineage, including both cytotoxic NK cells and non-cytotoxic ILC1s, necessitated a prompt reevaluation of their unique contributions to immunity and pathology. However, the inherent similarity^25,78^ and plasticity between NK cells and ILC1s^19–24^ obstructed their precise delineation and the elucidation of their unique contributions to immune regulation. This study introduces an innovative paradigm, exploiting the unique association of AsGM1 with cytotoxic attributes^44–48^—absent in ILC1s^34,37^—as a definitive criterion to distinguish ILC1s from NK cells. By prioritizing cytotoxic potential as the cardinal differentiator, our AsGM1-centric approach achieved accurate delineation, unveiling a precise view of the Group 1 ILC landscape and its remarkable plasticity across the health-disease continuum. An ideal demarcation marker is *i)* surface bound to preserve live cell integrity and functional potential, *ii)* exhibits consistency across tissues and physiological contexts, and *iii)* facilitates precise subset separation to minimize cell subset contamination risk. Against these benchmarks, AsGM1 emerged as a superior demarcating marker for Group 1 ILCs, surpassing traditional markers in precision and reliability. Its ability to distinguish ILC1s from NK cells across a spectrum of tissues - lymphoid, non-lymphoid, and mucosal was validated, with functional assays demonstrating its accuracy in identifying cytotoxic versus non-cytotoxic cellular identities within the Group 1 ILC lineage. RNA-seq analysis of AsGM1^-^ and AsGM1^+^ Group 1 ILCs confirmed their distinct identities as authentic ILC1s and NK cells, respectively. Corroborated by existing transcriptomic datasets, our findings established the efficacy of AsGM1 in segregating ILC1 and NK cell populations with equivalent accuracy to traditional multi-marker strategies^37,57^. Likewise, extended analysis incorporating existing multi-tissue datasets^37,57^ revealed a uniform core transcriptomic signature associated with AsGM1, underscoring its potential as a versatile, site agnostic marker for the systematic delineation of NK cells and ILC1s. This capability extends beyond steady state classifications, adeptly capturing the binary classification of NK cells and ILC1s during immediate immune responses, as demonstrated by our findings in the acute liver injury model. However, the landscape of cellular identity within the Group 1 ILC pool becomes markedly more complex in chronic infection scenarios, such as those driven by *Toxoplasma gondii*^23^, or within specific homeostatic contexts like the salivary glands^19^. In these cases, the binary classification of NK cells and ILC1s is disrupted by intra-lineage plasticity, namely NK-to-ILC1 plasticity, leading to the generation of transitional states that blur the boundaries between NK cells and ILC1s, resulting in a continuum of identities^19,23^. Remarkably, AsGM1 performance across these conditions revealed an unprecedented capability to trace NK-to-ILC1 plasticity, adeptly transcending the current challenges in distinguishing stable from transitional states.

NK-to-ILC1 plasticity represents a pivotal mechanism of adaptation characterized by the cellular reprogramming of NK cells into entities exhibiting ILC1-like attributes^19–25^. It involves the downregulation of NK cell markers (Eomes or CD49b) and upregulation of ILC1 markers (CD200R, CD49a, CD103, or TRAIL), resulting in a significant reduction of cytotoxic capabilities. These reprogrammed NK cells, now closely resembling ILC1s, shift their functional priority towards maintaining tissue integrity over executing direct cytotoxic actions, underscoring the formidable challenge of discerning the true identity of ILC1-like entities—whether they are authentic ILC1s or the products of NK cell plasticity. Validated in both TGFβ-driven^19^ and infection-driven^23^ models of NK-to-ILC1 plasticity, our findings highlight AsGM1 as a pivotal marker for tracking these cellular transitions, unaffected by the downregulation of Eomes or the upregulation of CD200R, CD49a, CD103, or TRAIL. Building upon the TGFβ-driven NK-to-ILC1 plasticity tracked via AsGM1 expression from NK cells (AsGM1^+^) to SG ILC (AsGM1^+^) in adult SGs^19^, analysis of aged tissues uncovered a distinct, age-related SG ILC1-like subset characterized by AsGM1^+^Eomes^-^ CD200R^+^CD49b^-^CD49a^+^ profile, suggesting additional layers of plasticity driven by aging. This age-dependent subset closely mirrors classical ILC1 profile, nearly to the point of potential misidentification. However, an unorthodox expression of AsGM1 at this juncture rather points to an advanced state of plasticity within the NK-to-ILC1 continuum^20^, likely influenced by the specific aging microenvironment which is characterized by substantial shifts in epigenetic landscapes^79^ and cytokine profiles^80^.

Leveraging insights from our investigation into tracing NK-to-ILC1 plasticity during aging, scRNA-seq analysis of SG Group 1 ILCs compellingly supports and further extends our findings at a transcriptomic level^56^. This analysis delineated five clusters, including NK cells, two ILC1 clusters, and two transitional states, aligning with TGFβ-driven plasticity in adult SGs^56^. Notably, PCA did not segregate these clusters; rather, it unveiled a continuum of cellular states. Further, Pseudotime analysis traced these clusters from NK cells to ILC1s via transitional stages, marked by the sequential downregulation of NK cell core markers and upregulation of ILC1-related genes. This trajectory not only corroborates the plasticity continuum we have mapped through AsGM1 tracing in aged SGs—from AsGM1^+^ NK cells to AsGM1^+^ SG ILCs and onto AsGM1^+^ SG ILC1-like cells— but also reveals the inherent potential for such transitions in adult tissues, a potential that becomes markedly manifest with age. Subsequent analysis of NK-to-ILC1 plasticity driven by *T. gondii* infection^23^ identified a unique AsGM1 expression gradient, sharply distinguishing this infection-driven plasticity from homeostatic-driven changes in the SGs. This gradient facilitated the precise identification of two ILC1-like states through intermediate AsGM1 expression, uncovering a previously characterized ILC1-like subset primarily observed in infected livers^23^, and revealing a distinct ILC1-like subset lacking NCR1, predominantly observed in infected spleens and lungs. The downregulation of NCR1 on NK cells, a phenotype prevalent in cancer^81–88^, and its association with impaired cytotoxicity and increased IFNγ production^89^, likely reflects a strategic adaptation to *T. gondii* infection, prioritizing tissue integrity and control of inflammation over cytotoxic actions. These four distinct ILC1-like entities captured by AsGM1 expression in contexts of aging and infection illustrate the expanding universe of Group 1 ILCs which strategically utilize NK-to-ILC1 plasticity to finely tune the equilibrium between pathogen defense and tissue homeostasis as needed.

The central point of this study lies in the resilience of AsGM1—a membrane lipid marker^41–43^—sharply contrasting with the loss of Eomes during NK-to-ILC1 plasticity. This persistent expression of AsGM1 in ILC1-like entities, despite cellular reprogramming and shifts in protein expression, is attributed to the inherent characteristics of lipids, which are less affected by transcriptional changes that directly influence protein expression^90,91^. The stability of lipids, crucial for membrane structure, is maintained through mechanisms like enzymatic regulation and metabolic processes, rather than direct transcriptional control^90^. Specifically, AsGM1 regulation through glycosyltransferases, rather than gene expression changes, underscores a distinct pathway of control^91^. Consistent with its established link to cytotoxic functions^44–48^, AsGM1 was traced to the immature stage of the NK cell developmental pathway, highlighting its early involvement at a crucial point of functional commitment^49,92^. This early expression together with its proven stability, unaffected by the transcriptional reprogramming that alters Eomes, establishes AsGM1 as an unparalleled marker for fate mapping NK-to-ILC1 plasticity. The prevalence of intra-lineage plasticity as a mechanism of adaptation across diverse contexts and species^9–24^, leading to distinct transitional entities, underscores the importance of our findings and the value of AsGM1 tracing for the accurate classification of entities within the NK-to-ILC1 spectrum as either stable or in a state of plasticity.

## METHODS

### Mice

C57BL/6 and *Rag2^-/-^* mice were purchased from Jackson Laboratories and maintained at the University of Michigan. CD11c^dnR^ transgenic mice^75,76^ were maintained on C57BL/6 background at the University of Michigan. Adult mice aged between 6 to 15 weeks, as well as aged mice between 30 to 52 weeks, were used in this study. All mice were maintained at the University of Michigan in specific pathogen-free housing. The University of Michigan approved the use of rodents through the criteria in the Guide for the Care and Use of Laboratory Animals from the National Institutes of Health. All procedures described in this study were approved by IACUC.

### Carbon Tetrachloride (CCl_4_) Acute Liver Injury

Adult C57BL/6 mice were injected intraperitoneally with 10μl of 10% CCl_4_ (02671; Sigma) diluted in peanut oil (P2144; Sigma) per gram of body weight^73^. Eighteen hours post-treatment, livers were perfused with cold PBS and processed for analysis.

### Parasitic Infection

*T. gondii* ME49 strain tachyzoites were cultured in primary human foreskin fibroblasts (HFF) in Dulbecco’s modified Eagle’s medium (DMEM) supplemented with 10% Cosmic Calf serum (Hyclone), 20 mM HEPES, 5 µg/mL penicillin/ streptomycin and 2 mM L-glutamine. Infected monolayers were scraped, ruptured to release the parasites by passage through a syringe with a 22g needle, and tachyzoites were recovered after passage through a 3 µm filter in PBS and one wash in PBS. Adult female C57BL/6 mice were injected intraperitoneally with 200 ME49 strain tachyzoites in PBS^23^. Mice were euthanized 14-15 days post infection and perfused with cold PBS. Spleens, lungs, and livers were processed for analysis.

### Organ Processing

Livers were harvested after perfusion with cold PBS and digested in 5% FBS medium (RPMI) supplemented with 0.01% collagenase IV (Sigma) and 0.001% DNase I (Sigma) for 30 min at 37°C under agitation (250 rpm). Liver mononuclear cells were obtained using Percoll gradient (Thermo Fisher) at the interface between 33% and 70% layers. Lungs were digested in 15ml total of 10% FBS medium (RPMI) supplemented with 1 mg/ml collagenase IV and 0.5 mg/ml DNase I for 30 min at 37°C under agitation (250 rpm).

Salivary (sublingual and submandibular) glands were digested in 10ml total of 5% FBS medium (RPMI) supplemented with 1ul HEPES, 5mM CaCl2, 0.5 mg/ml collagenase IV and 0.1 mg/ml DNase I for 60 min at 37°C under agitation (250 rpm). Salivary gland mononuclear cells were obtained using Percoll gradient at the interface between 40% and 70% layers^19^. To isolate intestinal cells, small intestine segments were first removed, and Peyer’s patches excised. After discarding and washing gut content, the small intestines were cut into small pieces and incubated in HBSS supplemented with 5% FBS, 2 mM EDTA (Fisher Scientific), 1 mM DTT (American Bioanalytical), and 10 μg/ml Gentamicin (Gibco) for 20 min at 37°C under agitation (250 rpm) to dissociate epithelial cells. Intraepithelial lymphocytes (IEL) were isolated from the supernatants by filtering through nylon wool (Polysciences) and discontinuous Percoll (GE Life Sciences) gradient at the interface between 33% and 70% layers. After a step of washing, intestinal pieces were digested in 5% FBS medium (RPMI) supplemented with 1 mg/ml collagenase D (Roche) and 0.5 mg/ml DNase I (Sigma) for 60 min at 37°C under agitation (250 rpm). Lamina propria mononuclear cells were obtained by Percoll gradient at the interface between 33% and 70% layers. To harvest PBMCs, blood was drawn by cardiac puncture after euthanasia and placed in a tube with EDTA to prevent clotting. Mononuclear cells obtained from livers, lungs, salivary glands, small intestine lamina propria, and IEL, as well as cells harvested from spleens, bone marrow, mesenteric lymph nodes, Peyer’s patches, and blood were filtered through 100μm cell strainer at final step before used for analysis. Red cells in the bone marrow, spleen, and blood were lysed using red cell lysis buffer (Invitrogen).

### In Vitro Stimulation

Cells from spleens and small intestine lamina propria were stimulated with 5 ng/ml IL-12 (Peprotech) plus 25 ng/ml IL-18 (R&D Systems) or 5 ng/ml IL-12 (Peprotech) plus 50 ng/ml IL-15 (Peprotech) for 4-16 hours at 37C with the protein-transporter inhibitor Golgi Stop (BD Biosciences) added during the last 4 hours of culture. Stimulated cells were analyzed for intracellular expression of IFNγ and GzmB. When indicated, cells from C57BL/6 and CD11c^dnR^ spleens were cultured for 7 days with 10 ng/ml IL-15 (PeproTech) supplemented with or without 10 ng/ml TGFβ1 (R&D). Cytokines were added on days 0 and 3, and cells were analyzed on day 7 using LIVE/DEAD™ Fixable Aqua Dead Cell Stain Kit (Invitrogen) to exclude dead cells.

### Flow Cytometry and Cell Sorting

Single-cell suspensions from the liver, lung, salivary glands, small intestine lamina propria, PPs, IEL, spleen, bone marrow, blood, and mesenteric lymph nodes were first treated with Fc receptor block (anti-CD16/CD32 clone 2.4G2) and then stained using the following fluorochrome-conjugated antibodies purchased from Biolegend, eBioscience, or BD Biosciences: anti-CD45 (30-F11), anti-CD3 (17A2), anti-NK1.1 (S17016D and PK136), anti-NCR1 (29A1.4), anti-AsGM1 (Poly21460), anti-CD200R (OX-110), anti-CD49b (DX5), anti-CD49a (HMa1), anti-CD103 (2E7), anti-CD25 (7D4), anti-Ly6C (HK1.4), anti-TRAIL (N2B2), anti-Ly49C/I (5E6), anti-Ly49H (3D10), anti-Ly49D (4E5), anti-Ly49G2 (4D11), anti-NKG2ACE (20d5), anti-KLRG1 (2F1/KLRG1), anti-CD94 (18d3), anti-T-bet (4B10), anti-Eomes (W17001A), anti-RORγt (B2D), anti-IFNγ (XMG1.2), anti-GzmB (NGZB), anti-CD122 (TM-BETA 1), anti-CD27 (LG.7F9), and anti-CD11b (M1/70). For intracellular staining, cells were fixed and permeabilized using FOXP3/Transcription factor fixation/permeabilization kit (Invitrogen). Stained cells were acquired on BD FACSCanto or BD LSRFortessa flow cytometers and analysis was performed using FlowJo software. For cell sorting, liver mononuclear cells were stained with anti-CD45, anti-NK1.1, anti-NCR1, and anti-AsGM1 antibodies. Group 1 ILCs were sorted based on AsGM1 expression as follows: AsGM1^+^ ILCs; CD45^+^NK1.1^+^NCR1^+^ AsGM1^+^ and AsGM1^-^ ILCs; CD45^+^NK1.1^+^NCR1^+^AsGM1^-^. Cells were sorted on FACSAria High-Speed Cell Sorter at the flow cytometry core facility (University of Michigan).

### RNA Preparation and Sequencing

Total RNA was isolated from AsGM1^+^ and AsGM1^-^ ILC subsets using the RNeasy Plus Mini Kit (Qiagen) according to the manufacturer’s procedure. RNA samples were treated with the DNA-free kit (Ambion) to remove any traces of genomic DNA. RNA was assessed for quality using the BioAnalyzer (Agilent #5067-1513). The SMART-Seq Stranded kit (Takara #634444) was used for cDNA synthesis and amplification, ZapR-mediated ribosomal RNA depletion, and library preparation from 1.8 ng of total RNA. Final libraries were checked for quality and quantity by Qubit hsDNA (Thermofisher #Q33231) and LabChip (Perkin Elmer #CLS1444006). The samples were pooled and sequenced on the Illumina NovaSeq S4 Paired-end 150bp, according to manufacturer’s protocols.

### RNA-Seq Analysis

The reads were trimmed using Cutadapt v2.3^93^. FastQC v0.11.8 was used to ensure the quality of data^94^. Fastq Screen v was used to screen for various types of contamination^95^.

Reads were mapped to the reference genome GRCm38 (ENSEMBL), using STAR v2.7.8a^96^ and assigned count estimates to genes with RSEM v1.3.3^97^. Alignment options followed ENCODE standards for RNA-seq. Multiqc v1.7 compiled the results from several of these tools and provided a detailed and comprehensive quality control report^98^. Data were pre-filtered to remove genes with less than 10 counts in total from all samples. Differential gene expression analysis was performed using DESeq2^99^ using a negative binomial generalized linear model (thresholds: log2 fold change >1 or <-1, Benjamini-Hochberg FDR (Padj) <0.05). Plots were generated using variations of DESeq2 plotting functions and dplyr^100^, pheatmap^101^, ggplot2^102^, ggrepel^103^, and ggbreak^104^ packages with R version 4.3.0.^105^ Annotation data from ENSEMBL 102 was used, and genes were additionally annotated with Entrez GeneIDs and text descriptions. Heatmaps and violin plots were generated from vst normalization values generated by DESeq2, and heatmaps were clustered by heatmap function using Euclidean distance and complete linkage. Public differentially expressed gene tables^37,57^ were compared to the differentially expressed genes found in our study. Differentially expressed genes from all three analyses were processed with Python to create a Pandas data frame with the Log2Fold change values of the common differentially expressed genes. The merged data frame was input into bioinfokit^106^ to generate the heatmaps. The Venn diagram was produced using VennDiagram based on the differentially expressed genes.

### Statistics

Prism 10 software (GraphPad) was used for all statistical analysis. Unpaired t-test, one-way ANOVA, or two-way ANOVA were used for the statistical analysis of differences between two or more groups with *p* < 0.05 considered significant and indicated by asterisks: \*\*\*\**p* < 0.0001, \*\*\**p* < 0.001, \*\**p* < 0.01, \**p* < 0.05. ns for non-significant *p* values.

## Acknowledgment

Library prep and next-generation sequencing were carried out in the Advanced Genomics Core at the University of Michigan. We acknowledge support from the Bioinformatics Core of the University of Michigan Medical School’s Biomedical Research Core Facilities (RRID:SCR_019168). Research reported in this publication was supported by the National Cancer Institute of the National Institutes of Health under award number P30CA046592 and the National Institute of Allergy and Infectious Diseases (R21 AI14010602 and R21 AI14010602; to Y.L. and X.S; and R01 AI120607 to V.B.C.).

