## Supplementary Figures 1-5 for "A Membrane Lipid Signature Unravels the Dynamic Landscape of Group 1 ILCs"

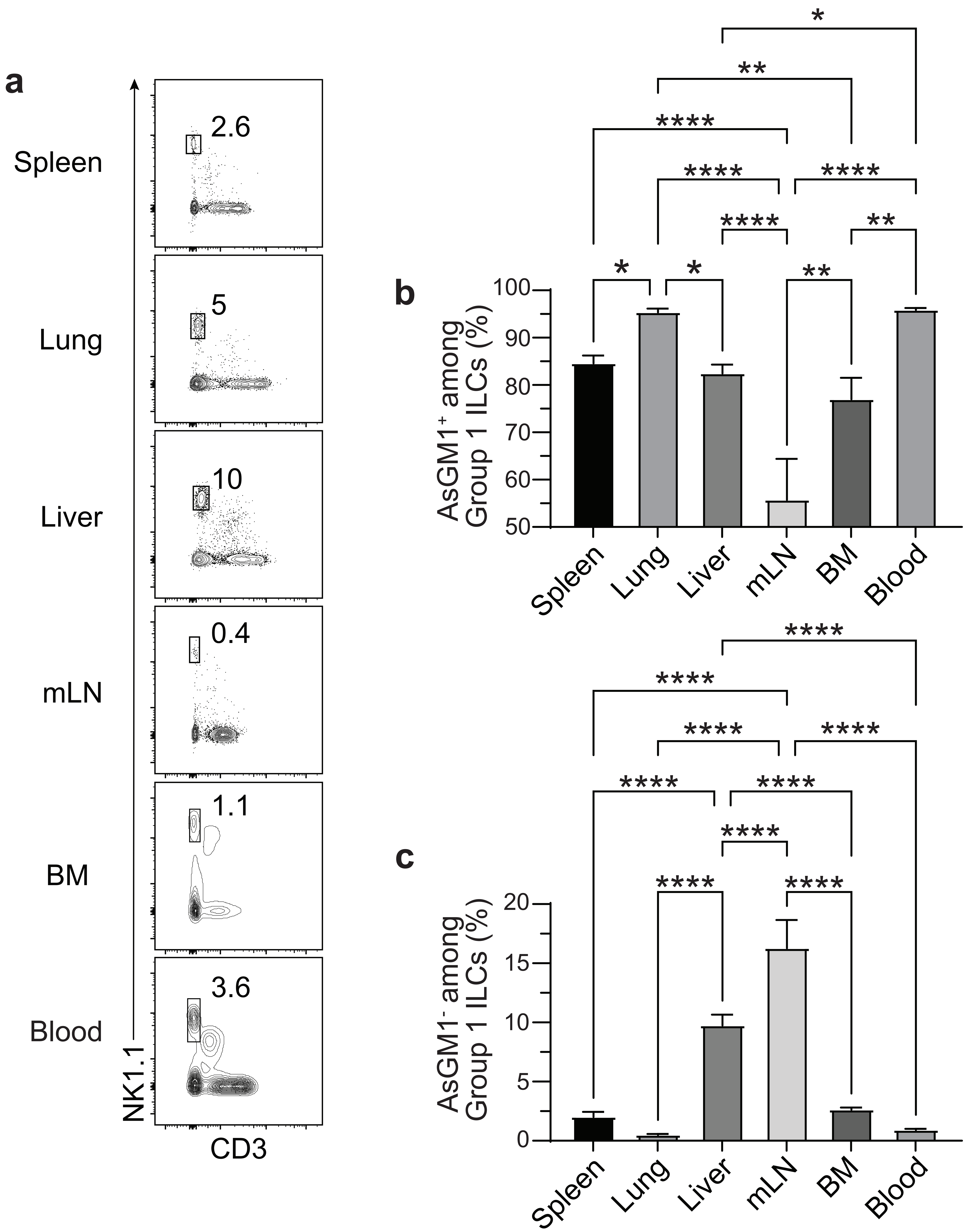


**Supplementary Figure 1. Distribution of AsGM1- and AsGM1+ Group 1 ILCs across tissues.** (**a**) Distribution of NK1.1 vs CD3 in the spleen, lung, liver, mesenteric lymph nodes (mLN), bone marrow (BM), and blood. (**b, c**) Frequency of $AsGM1^{+}$ (**b**) and $AsGM1^{-}$ (**c**) cells among Group 1 ILCs ($CD3^{-}NK1.1^{+}NCR1^{+}$) across tissues. Data represents 3 independent experiments: spleen (*n* = 27), lung (*n* = 5), liver (*n* = 16), mLN (*n* = 7), BM (*n* = 7), and blood (*n* = 7). Statistical validation employed one-way ANOVA (**b, c**), with significance indicated by *p<0.05, **p<0.01, ****p<0.0001. Error bars show mean ± s.e.m.

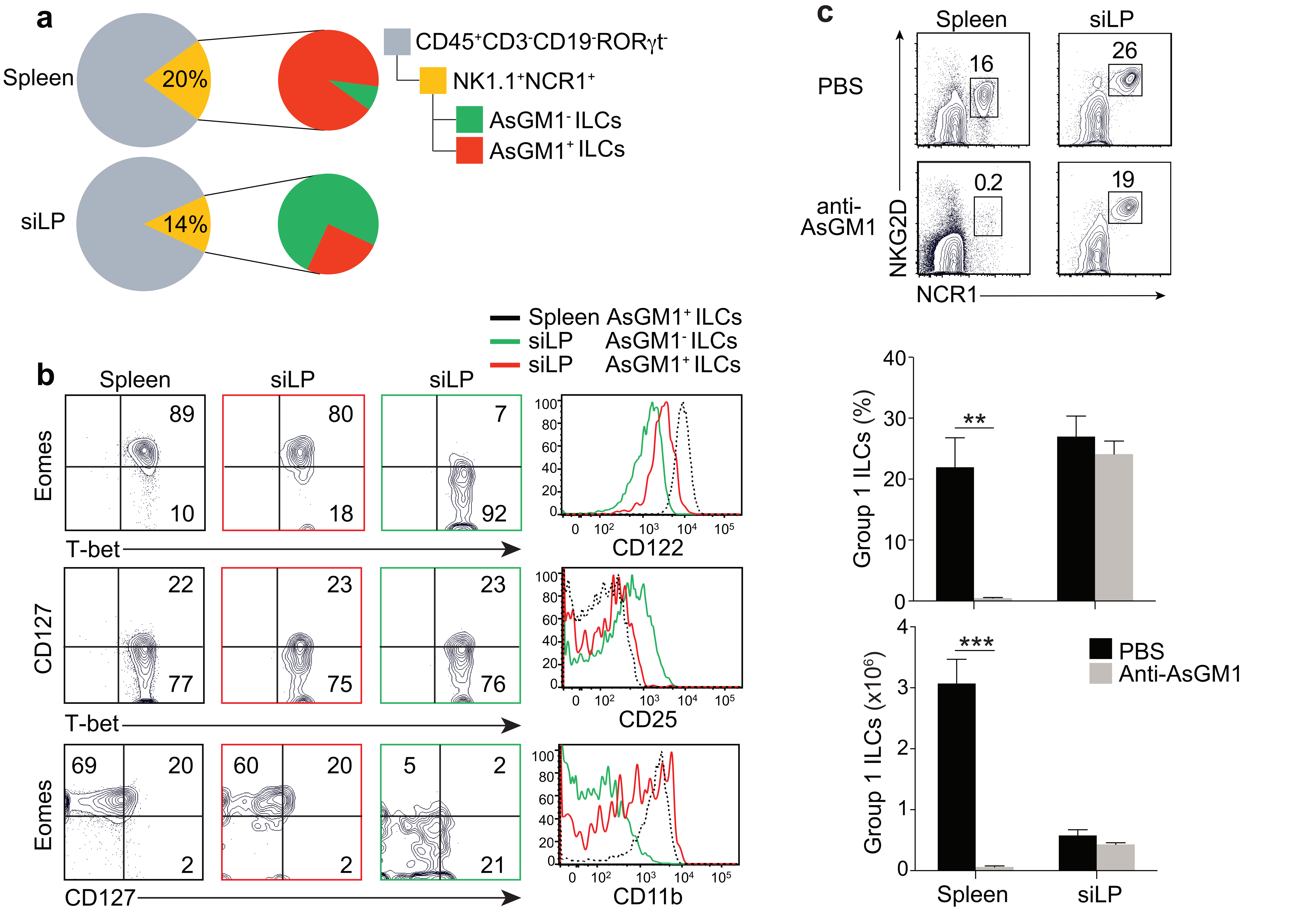


**Supplementary Figure 2. Differential effects of anti-AsGM1 treatment on splenic and intestinal Group 1 ILCs.**
(**a**) Ratio of AsGM1⁻ (green) to AsGM1⁺ (red) cells within Group 1 ILCs (yellow) in spleen and small intestine lamina propria (siLP). (**b**) Expression of Eomes, T-bet, CD127, CD122, CD25, and CD11b in AsGM1⁻ (green) and AsGM1⁺ (red) Group 1 ILCs in the siLP. Splenic AsGM1⁺ Group 1 ILCs (black) serve as reference. (**c**) Anti-Asialo-GM1 (Rabbit, FUJIFILM Wako) was diluted in 1 mL sterile water and 50 μl was administered *i.p.* into C57BL/6 mice on days 0 and 2. Control C57BL/6 mice received 2 doses of PBS on days 0 and 2. On day 3, mice were euthanized and Group 1 ILCs were analyzed in the spleen and small intestine lamina propria (siLP). Distribution of NKG2D and NCR1 among CD45⁺CD3⁻CD19⁻Rorγt⁻ cells. Frequency and numbers of Group 1 ILCs in the spleen and siLP of PBS (black) and anti-AsGM1 (grey) treated mice. Data represents 2 independent experiments with $n$ = 6 (**a, b**) and $n$ = 8 mice (**c**). Statistical validation employed two-way ANOVA, with significance indicated by **p<0.01, ***p<0.001. Error bars show mean ± s.e.m.

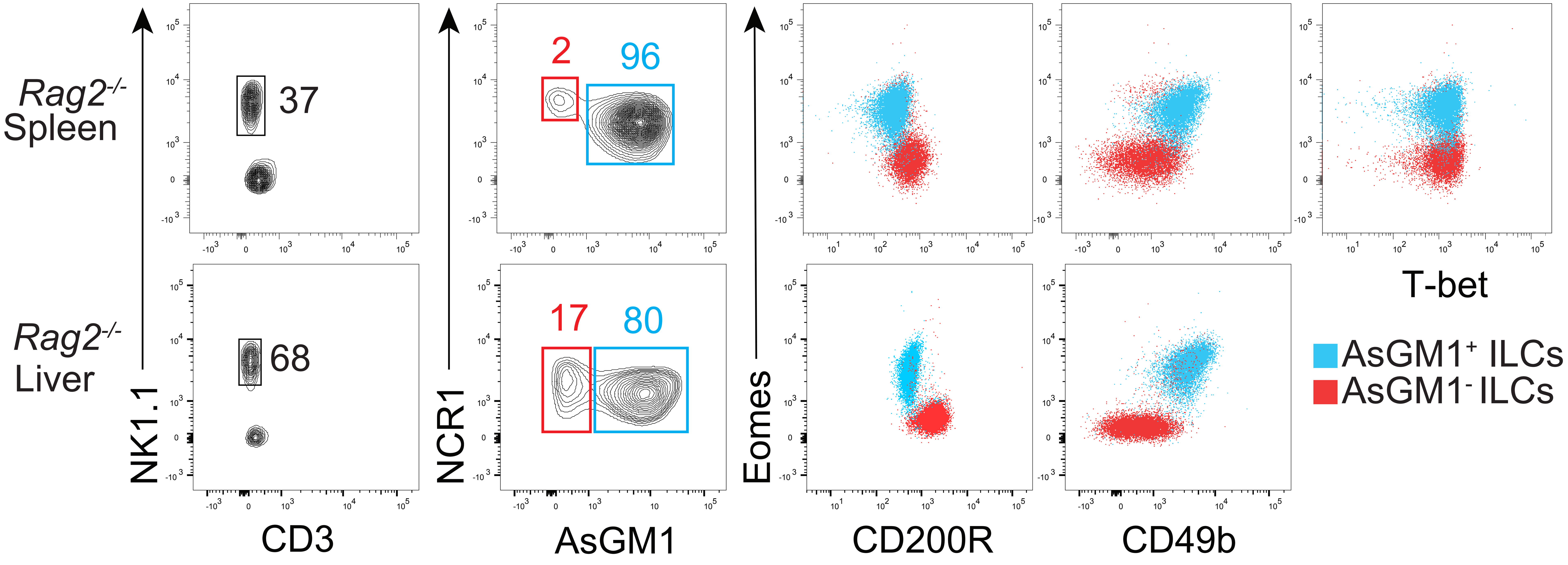


**Supplementary Figure 3. AsGM1$^-$ and AsGM1$^+$ Group 1 ILCs from *Rag2$^{-/-}$* mice used as input for RNA-seq analysis.** NK1.1 vs. CD3 distribution in the spleen and liver of *Rag2$^{-/-}$* mice. NCR1 vs. AsGM1 distribution among CD3$^-$NK1.1$^+$ ILCs. Distribution of Eomes vs. CD200R, CD49b, and T-bet among AsGM1$^-$ (red) and AsGM1$^+$ (blue) Group 1 ILCs. Data represent 3 independent experiments: spleen (*n* = 6) and liver (*n* = 4).

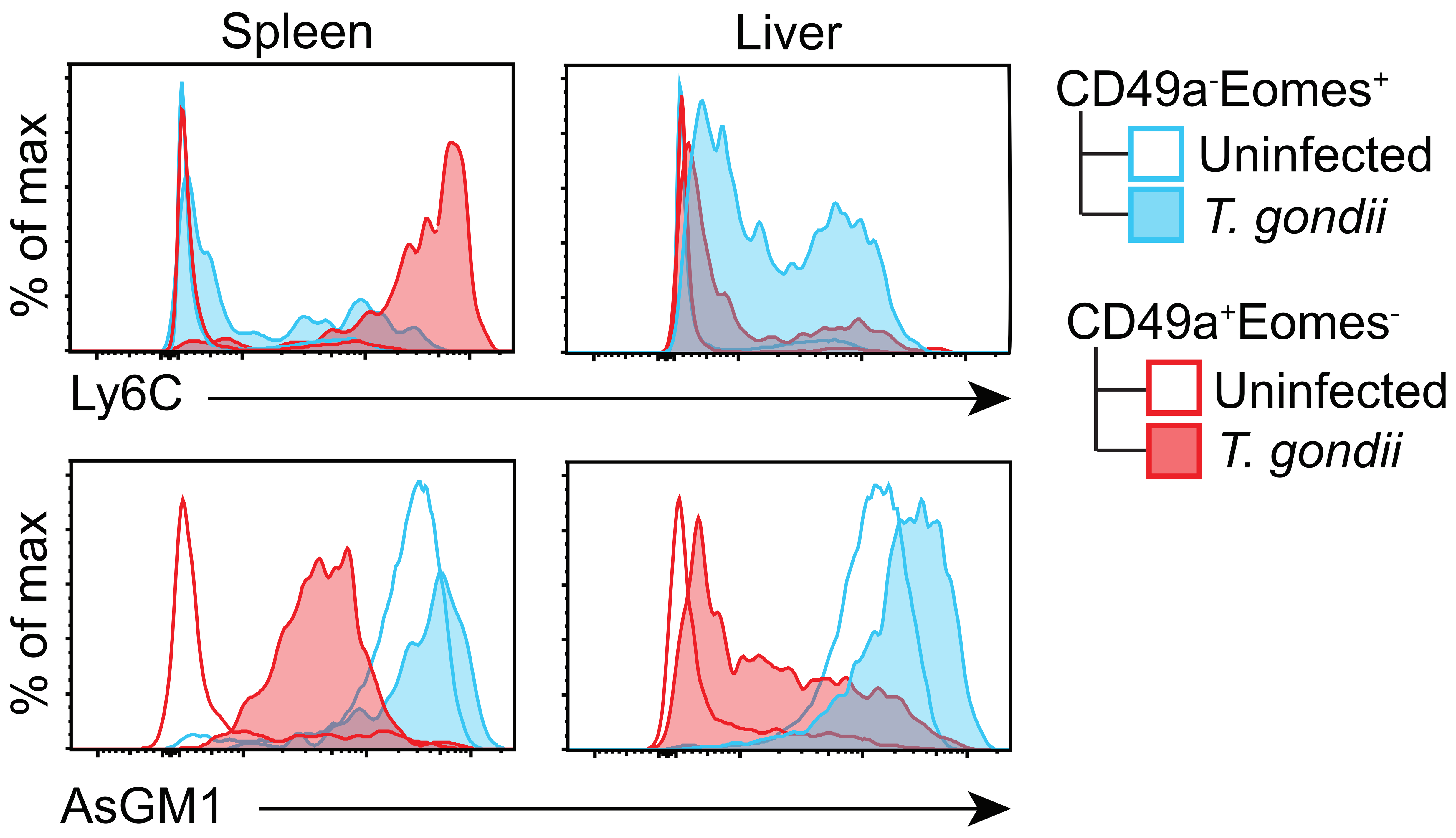


**Supplementary Figure 4. Alterations within the Group 1 ILC compartment driven by *Toxoplasma gondii* infection**. Expression of Ly6C and AsGM1 in ILC1s ($CD49a^+Eomes^-$) and NK cells ($CD49a^-Eomes^+$) from the spleen and liver of uninfected or *T. gondii* infected mice 14 days post-infection. Data represent 2 independent experiments, with $n$ = 6-11 uninfected mice and $n$ = 11-16 infected mice.

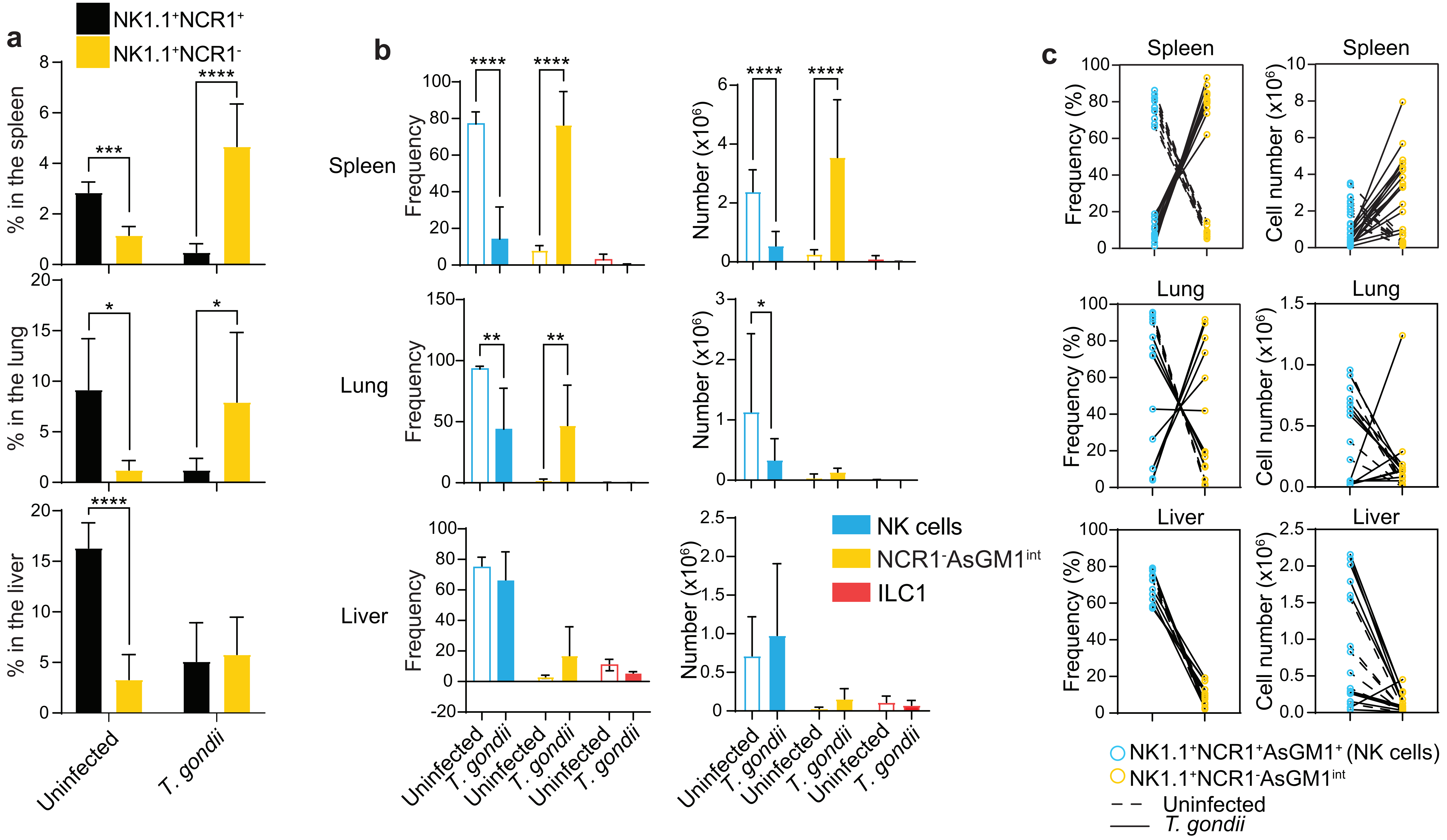


**Supplementary Figure 5. Greater diversity within Group 1 ILCs driven by *Toxoplasma gondii* infection.**
(**a**) Frequency of NCR1$^+$ and NCR1$^-$ Group 1 ILCs in the spleen, lung, and liver of uninfected or *T. gondii* infected mice 14 days post-infection. (**b**) Frequency NK cells (AsGM1$^+$NCR1$^+$; blue), NCR1$^-$AsGM1$^{int}$ ILC1-like cells (AsGM1$^{int}$NCR1$^-$; yellow), and ILC1s (AsGM1$^-$NCR1$^+$; red) among NK1.1$^+$ innate cells in the spleen, lung, and liver of uninfected or *T. gondii* infected mice 14 days post-infection. (**c**) Correlation between frequency and number of NK cells vs. NCR1$^-$AsGM1$^{int}$ ILCs in the spleen, lung, and liver of uninfected and *T. gondii* infected mice 14 days post infection. Each line represents an individual mouse. Data represent 2 independent experiments, with $n$ = 6-11 uninfected mice and $n$ = 11-16 infected mice. Statistical validation employed two-way ANOVA, with significance indicated by *$p$<0.05, **$p$<0.01, ****$p$<0.0001. Error bars show mean $\pm$ s.d.
